# Sensing the rainbow: genetic and physiological responses to light quality in Ostreococcus, an ecologically important photosynthetic picoeukaryote

**DOI:** 10.1101/2023.03.27.534389

**Authors:** Elizabeth Sands, Sian Davies, Richard John Puxty, François-Yves Bouget, David John Scanlan, Isabelle Alice Carré

## Abstract

Phytoplankton is exposed to dramatic variations in light quality as it moves up and down the water column or encounters the presence of sediments in the water. We investigated the potential impact on *Ostreococcus,* a key marine photosynthetic picoeukaryote, by analysing changes in its transcriptome, pigment content and photophysiology after acclimation to monochromatic red, green or blue light. The clade B species RCC809, isolated from the deep Atlantic Ocean, responded to blue light by accelerating cell division at the expense of storage reserves, and by increasing the relative level of blue-light absorbing pigments. In contrast, it responded to red and green light by increasing its potential for photoprotection. In contrast, the clade A species OTTH0595, which originates from a shallow water environment, showed no difference in photosynthetic properties and minor differences in carotenoid contents between light qualities. These results demonstrate that light quality can have a major influence on the physiology of eukaryotic phytoplankton, and suggest that different light quality environments can drive selection for diverse patterns of responsiveness and environmental niche partitioning.

**Highlight:** We characterise the effects of light quality on the transcriptome and photophysiology of *Ostreococcus*, a photosynthetic picoeukaryote, and show that responses are distinct between two ecotypes originating from different environments.

## Introduction

Phytoplankton are a vital part of the marine ecosystem. As primary producers, they provide a source of energy for the food web. They cycle nutrients, form a sink for carbon, and are responsible for roughly half the oxygen released to the atmosphere globally (Kulk *et al*., 2020). Photosynthetic picoeukaryotes make a major contribution to these processes, being responsible for 25-44% of global carbon capture (Jardillier *et al*., 2010; Kirkham *et al*., 2013; Mullin, 2001; Rii *et al*., 2016). It is therefore important to understand how these organisms adapt to various ecological niches in the ocean, and to identify the environmental factors that influence their abundance. Here we investigate the effects of light quality, a factor which is generally under-recognised in the marine environment.

Whilst the spectrum of incident light must match the absorption spectrum of photopigments for efficient photosynthesis, light quality can also serve as an indicator of competition by other organisms and initiate changes in growth strategy. On land, the light environment is broadly composed of light across the visible spectrum. Land plants possess blue, red, far-red and UV photoreceptors which enable them to respond to the spectral light environment. While red and blue light both promote photomorphogenesis, a low red to far-red ratio signifies shading by other plants. This triggers the shade avoidance response, which includes elongation of stems and petioles, an increase in leaf area, as well as accelerated flowering (Franklin and Whitelam, 2005). Distinct factors influence light quality in oceanic systems, however (Bouman *et al*., 2000). While visible light in the PAR range reaches a maximum depth of around 200m, red, far-red and green light wavelengths are absorbed in the upper layers of the water column, so that only blue light reaches the deep euphotic zone. The light spectrum is also affected by the presence of particulates in coastal waters. Light absorption and scattering by dissolved organic matter and run-off from rivers causes an increase in yellow wavelengths (Erga *et al*., 2012), while abundant phytoplankton increases the proportion of green light (Kume *et al*., 2018; Lichtenthaler, 1987; Schubert *et al*., 2001; Sun *et al*., 2010; Tilzer *et al*., 1995). Some species of cyanobacteria respond to these changes in light quality through a process known as complementary chromatic adaptation, during which levels of different phycobiliprotein pigments associated with the phycobilisome are adjusted to optimise absorption of excitation energy present in the environment [25-27]. Much less is known, however, about how eukaryotic microalgae respond to light quality in their environment.

There is mounting evidence that they do, however. Light quality-dependent changes in photopigments were observed in diatoms, suggesting a form of chromatic adaptation (Brunet *et al*., 2014; Hintz *et al*., 2021; Xu *et al*., 2021). Light quality was found to affect the protein content of the red alga *Porphyra leucosticte* [28], the diatom *Cyclotella nana* and the green alga *Dunaliella tertiolecta* [29], and the accumulation of lipids in the chlorophyte *Chlorella* sp. and the heterokonts *Nannochloropsis oculata and Nannochloropsis gaditana* (Patelou *et al*., 2020; Yuan *et al*., 2020). Light quality also impacted on growth rates of phytoplankton. Most species tested grew the fastest under blue *light* and the slowest under green light (Neun *et al*., 2022; Patelou *et al*., 2020; Yuan *et al*., 2020), and blue light promoted the highest rate of growth in controlled mesocosm experiments with natural phytoplankton communities (Hintz *et al*., 2021; Xu *et al*., 2021). *Dunaliella salina* showed the fastest growth rates under red light, however (Li *et al*., 2020), suggesting that distinct light quality responses may enable different species or ecotypes of microalgae to thrive in distinct environments, and therefore to occupy different ecological niches.

Here, the investigate the effects of light quality on the transcriptome, pigment content and photophysiology *of Ostreococcus,* a photosynthetic picoeukaryote found in oceans across the world, including tropical and temperate environments (Limardo *et al*., 2017; Rii *et al*., 2016). We show that two ecotypes originating from distinct light quality environments exhibit contrasting responses. These findings suggest that the differential ability to respond to light quality signals may contribute to the specialisation of specific phytoplankton ecotypes to different environments.

## Materials and Methods

### Cell lines and growth conditions

*Ostreococcus* ecotypes RCC809 (a clone of RCC141) and OTTH595 (also known as RCC745, RCC4221, OTTH 0595 or O. *tauri*) were obtained from the Roscoff Culture Collection (http://roscoff-culture-collection.org). Cells were grown in Keller medium (Keller *et al*., 1987). Batch cultures were incubated at 21°C under diurnal light-dark cycles composed of 12h light and 12h darkness (12L12D) from cool-white fluorescent bulbs at an intensity of 20μmol photons m^2^ s^1^. They were transferred to fresh medium every 14 days.

### Monochromatic light sources

Red and blue light was provided by LED arrays, and green light was provided by cool-white fluorescent bulbs covered with one layer of green filter (Lee filter 139). Light intensity was the same under all conditions, i.e. 4μmol photons m^-2^ s^-1^. Emission spectra from these different light sources are shown in Fig. S1.

### Determination of growth rates

Growth was monitored by measuring absorbance at 550nm, and absorbance was converted to cell number using the equation cell abundance (cells ml^-1^) = 2×10^9^ OD_550 nm_.

### Determination of cell size

Flow cytometry measurements of forward scatter signal height (FSC-H) were carried as described previously (Chretiennot-Dinet *et al*., 1995; Mullaney *et al*., 1969), 72 hours after transfer to monochromatic light. Fluoresbrite Multifluorescent beads with an average diameter of 0.5μm were used for calibration to allow comparison of relative cell size between light conditions.

### RNA-Seq experiment

Cells were grown in 200ml cultures in 1l flasks under white light as above until they reached mid to late log phase (to approximately 40 million cells per ml). They were then transferred to constant monochromatic red, green, or blue light. After 72 hours, cells were centrifuged at 5,000 x g, pooled into 1ml cold PBS, then spun again at maximum speed in a microfuge. The supernatant was discarded and the cell pellet frozen in liquid nitrogen. For RNA extraction, 1ml of TRIzol reagent was added to the frozen samples before thawing at room temperature. Two glass beads were added before shaking for 3 minutes using a Tissue Lyser (QIAGEN) at maximum speed. 200μl chloroform was added and mixed by shaking for 15 seconds before incubation at room temperature for 3 minutes and centrifugation at 12,000g at 4°C for 15 minutes. The aqueous phase was transferred to a fresh tube, combined with 0.5ml isopropanol and 5μl of glycogen (20mg/ml), incubated for 10 minutes at room temperature then spun down at 12,000g at 4°C for 15 minutes. The supernatant was removed, and the pellet washed twice with 1ml 70% (v/v) ethanol before resuspension in 50μl of RNAse free water. Samples were treated with RNAse-free DNAse (Sigma-Aldrich) according to the manufacturer’s instructions, before RNA purification using the Spectrum Plant Total RNA Kit (Sigma-Aldrich). RNA quality was verified using a Bioanalyser before preparation of RNA-Seq libraries using the Illumina Tru-Seq RNA library preparation kit, which enriches for mRNAs using Oligo-dT beads to capture polyA tails before cDNA synthesis. This method does not capture chloroplast RNA. 100bp paired-end sequencing was carried out on an Illumina HiSeq at the Wellcome Trust.

RNA-Seq data was analysed within the Cyverse Discovery Environment (https://www.cyverse.org). Contaminating adapter sequences and poor quality sequences were removed using Trimmomatic v 0.36.0 (Bolger *et al*., 2014). The Fastqc tool version 0.2 was then used to produce fastq files (Andrews, 2015), and TopHat version 2 to map reads to reference genomes using Bowtie 2 (Kim *et al*., 2013; Langmead and Salzberg, 2012). Samples which gave low numbers of reads were removed from further analyses. CuffDiff version 2.2.1a was used to calculate differential expression values for each gene in each pair of samples (Trapnell *et al*., 2013). Read counts were normalised by transcript length and by the total number of fragments, using the geometric Fragments per Kilobase of Transcript per Million fragments mapped (FPKM) method. Differentially expressed genes were identified based on corrected p values (q-values) and a false discovery rate (FDR) less than 0.05. This gave pairwise comparisons in the form of expression levels sorted by log2 fold change between each pair of light conditions. Gene expression patterns were visualised using the CummeRbund package (Goff *et al*., 2020) and heatmaps were refined using the pheatmap package (Kolde, 2019). Reference genomes used were the ORCAE OTTH595 v2 genome ((Blanc-Mathieu *et al*., 2014; Derelle *et al*., 2006; Palenik *et al*., 2007) and the RCC809 v2 genome (Grigoriev *et al*., 2012). Reference genome sequence (fasta) files and annotation (gff3) files for each *Ostreococcus* ecotype were sourced from the Online Resource for Community Annotation of Eukaryotes (ORCAE) database (http://bioinformatics.psb.ugent.be/orcae/).

### Gene Ontology (GO) enrichment analyses and KEGG pathway mapping

OTTH595 genes were annotated by compiling existing gene descriptions and GO terms from ORCAE and the Universal Protein knowledgebase (UniProt). In order to obtain GO-term annotations for RCC809 genes, a genome-wide BLAST search of the NCBI non-redundant database was carried out using the Diamond BLASTx command line tool: ‘translated Query-Protein Subject BLAST 2.2.31+’ (E value 1.00E-03) (Camacho *et al*., 2009). GO terms were then obtained using BLAST2GO software (Götz *et al*., 2008). These annotations were compared to and merged with those available from the ORCAE and Interpro databases and with annotations from OTTH595 homologues, identified by reciprocal BLAST. These were found to be consistent and, in some instances, more detailed than the existing annotations. GO-term over-representation analyses were carried out using the BiNGO 3.0.3 app in Cytoscape (Maere *et al*., 2005). Multiple testing correction was carried out using Benjamini & Hochberg False Discovery Rate (FDR) correction with a significance cut-off at 0.05.

Pathway mapping using KEGG was carried out by searching for the homologues of the RCC809 enzymes in OTTH0595 as RCC809 is not yet listed on KEGG. These homologues were identified using a reciprocal genome-wide BLAST based on a minimum of 50% identity.

### Promoter motif identification

Analysis of the intergenic region length distribution in RCC809 showed a peak around 200bp. This informed the selection of 250 base pair regions upstream of start codons for promoter analyses. Coordinates of these regions were collected in bed files, listing the genomic coordinates for these promoter sequences. Fasta sequence files were then generated from the bed files using the GetFastaBed tool in Galaxy (usegalaxy.org) (Gruening, 2014; Quinlan and Hall, 2010; The_Galaxy_Community, 2022). Short motifs that were over-represented in the promoters of light-responsive genes compared to a background of promoters genome-wide were identified using the Dreme 5.0.4 tool in the Meme suite (http://meme-suite.org) (Bailey, 2011). Matches to known or predicted binding sites for RCC809 transcription factors (TFs) were identified from the CIS-BP database using Motif Scan (Weirauch *et al*., 2014).

### Analysis of photosynthetic parameters

Cell cultures were grown under white light then acclimated to monochromatic light for 72 hours, before analysis of photosynthesis parameters using a PhytoPAM fluorometer. Cultures were diluted 5-fold before analysis to avoid saturation of the fluorescence signal. In order to determine the maximum quantum yield of photosystem II (PSII) photochemistry (Fv/Fm), samples were kept in darkness in the cuvette for 5 minutes, so that the primary electron acceptor would be fully oxidised and a basal fluorescence level (F0) could be measured. A saturating pulse of 470 nm light at maximum intensity (2600 µmol photons m^-2^ s^-1^) was then applied at 500ms intervals and maximum fluorescence (Fm) was measured. To determine the effective quantum yield of PSII photochemistry ( ), samples were subjected to increasing light intensities at 0, 3, Φ 6, 36, 94, 124, 184, 213, 270, and 298μmol photons m^-2^s^-1^ at 120s intervals so that a fluorescence steady state Ft would be reached. ΦPSII was calculated as (Fm’-Ft/Fm’) using the Phyto-Win software (V 2.13). The Phyto-Win software also calculated the relative electron transport rate (rETR) as Yield x PAR x 0.5 x 0.84μ the Platt curve fitting model to provide alpha values and IK values from rapid light curve data (Platt *et al*., 1980).

### Pigment analyses

Cell cultures were grown under white light, then acclimated to monochromatic light for 72 hours as for the RNA-Seq experiment. Cell cultures were then harvested onto GF/F Glass Microfiber filters (Whatman) using a vacuum pump. The filters were immediately flash frozen in cryovials in liquid nitrogen and stored at -80°C until shipped to the Danish Hydraulic Institute (DHI) laboratory in Denmark for analysis by high performance liquid chromatography (HPLC) as described by (Van Heukelem and Thomas, 2001). Pigment standards were not available for uriolide, micromonal, dihydrolutein, and for a previously described unknown carotenoid (Guyon *et al*., 2018) , so these were determined using the response factor for β-carotene. The abundance of two chlorophyll b-like pigments in RCC809 was calculated using the chlorophyll b response factor.

## Results

### Blue light accelerates cell division in the RCC809 ecotype

The RCC809 ecotype of *Ostreococcus* was first isolated from a deep ocean environment. In nature, it is exposed to a range of light conditions, ranging from bright, broad spectrum light near the surface, to dim blue light near the bottom of the euphotic zone. While its responses to light intensity have been described previously (Six *et al*., 2008; Six *et al*., 2009), effects of light quality have not been investigated.

To test for its ability to grow under different qualities of monochromatic light, RCC8909 cultures were grown under white light until cell densities reached 4 x 10^6^ cells.ml^-1^, then transferred to red, green, or blue light of equivalent photosynthetically active radiation (PAR) of 4μmol photons m^-2^ s^-1^. RCC809 cultures grew the fastest under blue light, reaching cell densities almost twice higher than under red and green light by the end of the experiment (**Fig. 1a**).

**Fig. 1.**
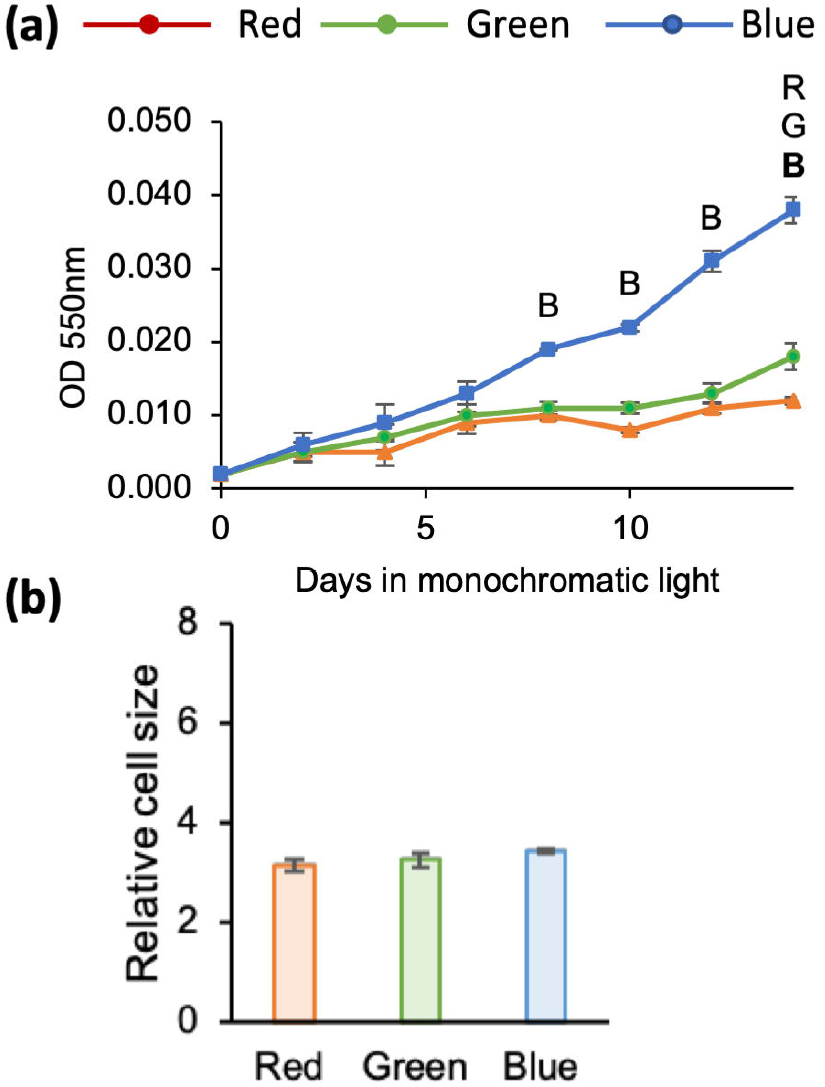
Effects of light quality on RCC809 cultures. **(a)** Growth kinetics. **(b)** Cell sizes estimated by FSC-H relative to singlet beads. Cultures were grown under red, blue or green monochromatic light at 4μmol m^-2^ s^-1^. Letters indicate significant differences between red (R), blue (B) or green (G) light (p < 0.05). Bold letters indicate p < 0.01.

To test whether the increased biomass under blue light might reflect changes in cell size rather than cell number, flow cytometry measurements of forward scatter signal height (FSC-H) were carried out. FSC-H was previously shown to positively correlate with cell size, and used to assess differences between *Ostreococcus* lineages (Mullaney *et al*., 1969; Schaum *et al*., 2016). However, no differences were observed between the different light conditions 72 hours after transfer to red, blue or green light (**Fig. 1b**), indicating that differences in biomass were solely linked to different rates of cell division.

### Transcriptional responses to light quality in RCC809

To gain an insight into which biological processes were affected by light quality, we compared the transcriptomes of cultures that were acclimated to constant, monochromatic red, green or blue light for 72 hours (**Fig. 2a**). Differences in gene expression were analysed by RNA-Seq. Genes that were differentially expressed in pairwise comparisons between light qualities were identified based on corrected p values and a false discovery rate (FDR) less than 0.05 (Table 1, Supplementary dataset 1). Principal component analysis (PCA) showed that samples collected under different light qualities clustered into different groups, consistent with differential gene expression (**Fig. 2b**). Blue light samples were the most distinct, whereas red and green samples were more similar to each other. Thus, the majority of differentially expressed genes had similar expression levels between red and green light conditions, but clearly different expression levels under blue light (**Fig. 2c**).

**Fig. 2.**
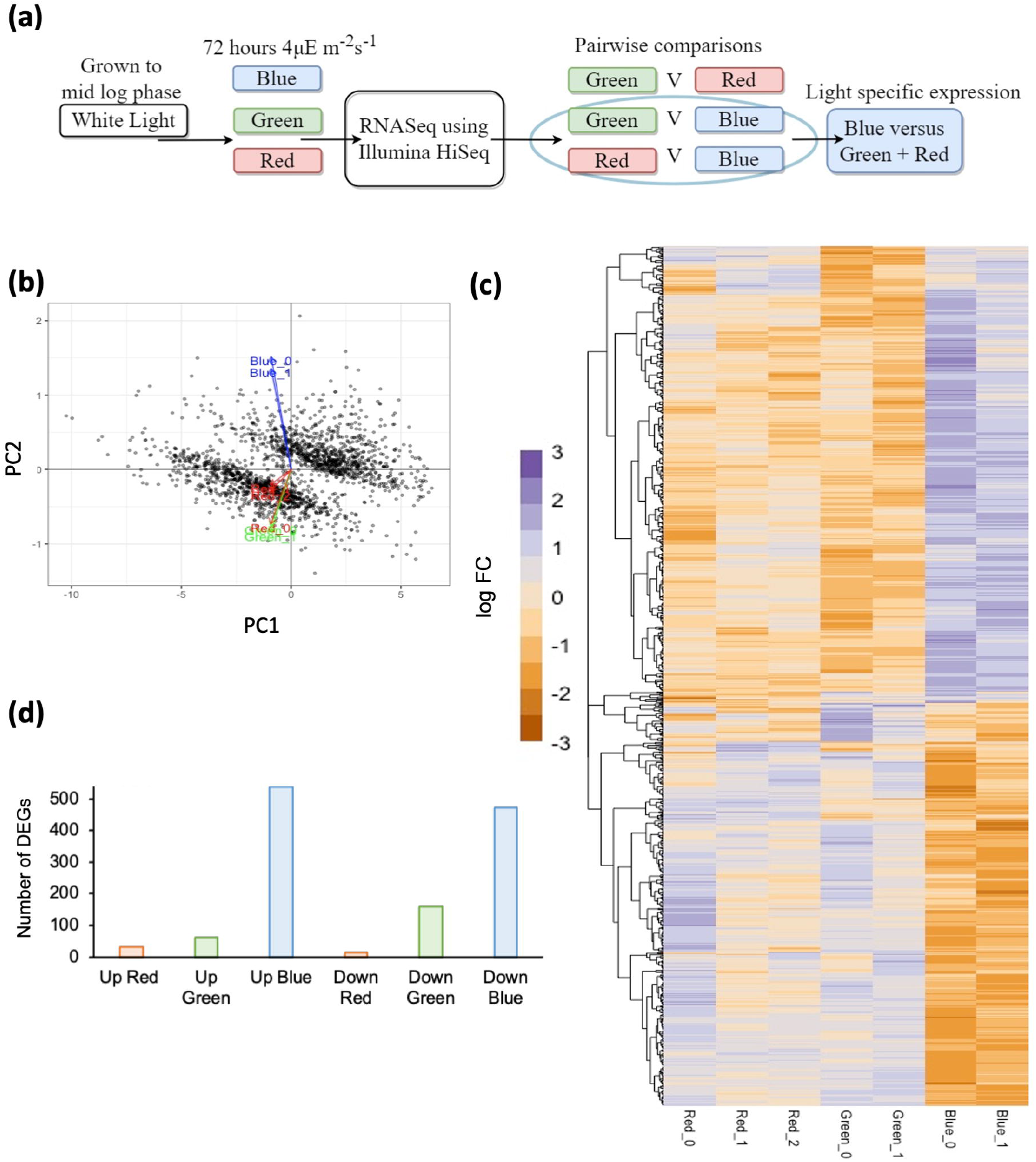
Transcriptomic analysis of RCC809 responses to light quality. **(a)** Experimental design. Cells were grown to mid-log phase then transferred to red, green or blue monochromatic light at 4μmol m^-2^ s^-1^ for 72 hours. Cultures were then sampled for RNA-Seq analyses. Light quality-responsive genes were initially identified through pairwise comparison of expression levels between light qualities. Responses specific to individual light qualities were then identified as differences in gene expression that were observed between one light quality and both others. **(b)** Principal component analysis of gene expression data. Red, green and blue arrows represent the variable vectors for individual samples under the corresponding light conditions. **(c)** Heatmap of differentially expressed genes (DEGs) identified from the RNA-seq analysis. Columns correspond to individual samples. Rows correspond to individual genes, clustered by Jensen-Shannon distance. Color from red to blue indicate log fold changes relative to mean expression levels. **(d)** Number of genes showing upregulation (Up) or downregulation (Down) specific to red, green or blue light.

**Table 1:** Identification of red, blue and green light-specific genes by RNA-seq. Differentially genes were first identified in pairwise comparisons between light conditions. Specific light quality responses were then identified as genes that were either up-regulated or down-regulated under one wavelength relative to both others.

| Differential expression | Number of DEGs | Light quality response | Number of light quality-responsive genes |
| --- | --- | --- | --- |
| Blue > Red | 706 | Up-Blue | 536 |
| Blue > Green | 807 | (Blue > Red AND Green) |  |
| Blue < Red | 661 | Down-Blue | 472 |
| Blue < Green | 746 | (Blue < Red AND Green) |  |
| Red > Blue | 666 | Up-Red | 32 |
| Red > Green | 237 | (Red > Blue AND Green) |  |
| Red < Blue | 706 | Down-Red | 14 |
| Red < Green | 113 | (Red < Blue AND Green) |  |
| Green > Blue | 746 | Up-Green | 32 |
| Green > Red | 113 | (Green > Blue AND Red) |  |
| Green < Blue | 807 | Down-Green | 159 |
| Green < Red | 237 | (Green < Blue AND Red) |  |

To separate gene expression responses by light quality, we identified genes whose expression was significantly increased or decreased under a given wavelength of light, compared to both others (Table 1). For example, genes that were induced by blue light were identified as significantly upregulated in blue light samples relative to both red and green light samples. This identified 536 genes that were specifically induced by blue light, and 472 that were repressed. On the other hand, only 32 genes were induced and 14 repressed by red light, whereas 60 were induced and 159 repressed by green light (Table 1, **Fig. 2d**).

Motif over-representation analyses were then carried out to identify candidate regulatory elements responsible for the different types of responses. Potential matches to transcription binding sites were identified from the cis-BP database (Table 2). Remarkably, the most frequent motifs within the promoter sequences of light quality-responsive genes matched binding sites for MYB transcriptions factors. The sequence GATATTT, found to be over-represented within genes that were down-regulated under red light, matched the known binding site for the MYB transcription factor CCA1 (Od04g00700), a light-responsive component of the *Ostreococcus* circadian clock (Corellou *et al*., 2009). The motif GGATAG, predicted to be bound by the MYB protein Od06g01160, was over-represented within genes that were induced by red light, as well as within genes that were repressed by blue light. On the other hand, the motif CGATTC, predicted to be recognised by the MYB transcription factor Od12g02060, was over-represented within genes that were upregulated under blue light. The binding site for the cell-cycle related transcription factor E2F (GTTCCCC) was also over-represented in this group, consistent with the increased rate of cell division observed under blue light, The data further suggested potential roles for the sequence CCACGTGG and a BHLH transcription factor (Od20g02280) to down-regulate gene expression in response to blue light.

**Table 2.**
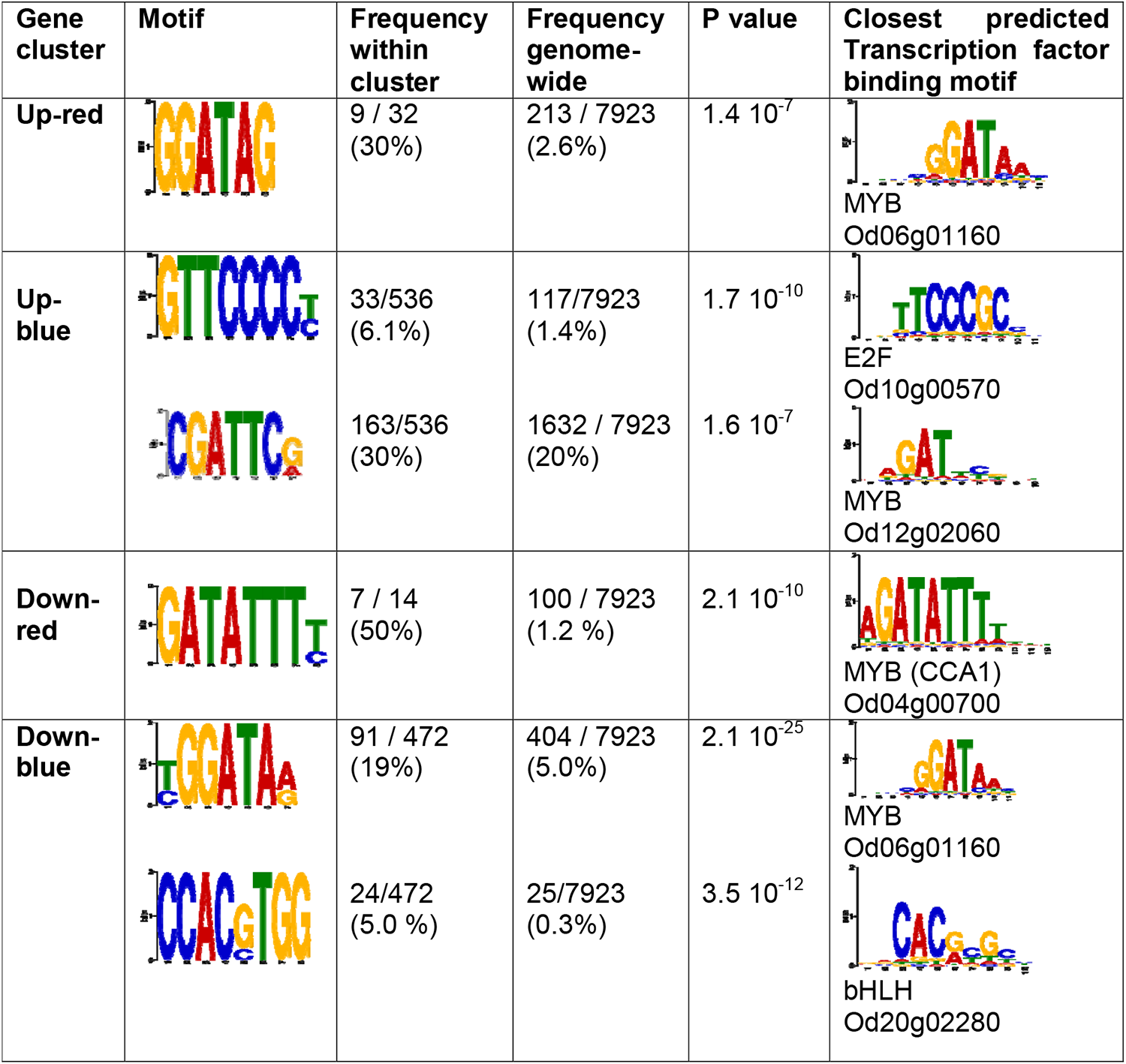
Putative regulatory motifs in the promoters of light quality-responsive genes. These were identified as over-represented in the 5’ upstream region of light quality-responsive genes relative to the whole genome. Cognate transcription factors were identified based on the closest match to a predicted transcription factor binding motif within the Cis-BP database 2.0 (Weirauch et al., 2014).

### Effects of light quality on biological processes in RCC809

Gene ontology (GO)-term analyses were carried out to test for enrichment of specific functional categories amongst light quality-responsive genes, relative to the rest of the genome. Differentially expressed genes were also mapped onto KEGG pathways to reveal potential metabolic consequences. This revealed effects of light quality on pathways related to cell division, primary metabolism, pigment biosynthesis and photosynthesis (Table 3).

**Table 3:** Overrepresented functional categories in RCC809 light quality-responsive genes.

| **GO: Biological Process** | **Corrected p-value** | **Number of genes (frequency)** | **Number genome-wide (frequency)** |
| --- | --- | --- | --- |
| **Downregulated in green light** |  |  |  |
| DNA replication | 5.75E-10 | 18 (18%) | 89 (2%) |
| DNA duplex unwinding | 1.08E-02 | 5 (4.9%) | 24 (0.5%) |
| Nitrate transport | 1.84E-02 | 2 (0.2%) | 2 (0.04%) |
| **Upregulated in red light** |  |  |  |
| NADPH regeneration | 2.56E-03 | 4 (16%) | 29 (0.7%) |
| Glycolysis | 1.29E-02 | 3 (12%) | 33 (0.8%) |
| Gluconeogenesis | 1.45E-02 | 3 (12%) | 37 (0.9%) |
| **Upregulated in blue light** |  |  |  |
| Cellular response to stress | 7.30E-03 | 29 (11%) | 204 (4.8%) |
| DNA-dependent DNA replication | 7.81E-03 | 11 (4.1%) | 42 (1.0%) |
| Chromosome segregation | 1.95E-02 | 7 (2.7%) | 21 (0.5%) |
| Tricarboxylic acid cycle | 2.20E-02 | 9 (3.4%) | 37 (0.9%) |
| **Downregulated in blue light** |  |  |  |
| Pigment biosynthetic process | 1.56E-07 | 18 (6.6%) | 47 (1.1%) |
| Chl biosynthetic process | 9.29E-07 | 11 (5.1%) | 19 (0.4%) |
| Glycolysis | 3.89E-03 | 9 (3.3%) | 33 (0.8%) |
| Gluconeogenesis | 7.91E-03 | 9 (3.3%) | 37 (0.9%) |
| Carotenoid (tetraterpenoid) biosynthetic process | 1.04E-02 | 5 (1.8%) | 12 (0.3%) |
| Nicotinamide nucleotide metabolic process | 1.49E-02 | 9 (3.2%) | 41 (1.0%) |
| Lipid biosynthetic process | 1.58E-02 | 18 (6.5%) | 126 (3.0%) |
| Photosynthesis | 2.45E-02 | 15 (5.5%) | 101 (2.4%) |
| Glucan biosynthetic process | 4.67E-02 | 4 (1.5%) | 11 (0.3%) |
| Reductive pentose-phosphate cycle | 4.67E-02 | 2(0.7%) | 2 (0.05%) |
| Endosome organization | 4.67E-02 | 2(0.7%) | 2 (0.05%) |
| Phylloquinone biosynthetic process | 4.67E-02 | 2(0.7%) | 2 (0.05%) |

41 genes related to the cell division cycle were differentially expressed under the different light qualities, including genes related to DNA replication, DNA repair and chromosome segregation (Table S1). Of these, 27 were upregulated under blue light relative to red and green light (Table S1). For example, the expression of cyclin-dependent kinase B (Od14g01080), which is upregulated during S-phase in *Ostreococcus* (Corellou *et al*., 2005; Farinas *et al*., 2006), more than doubled under blue light (Fig. S2), and expression of two homologues (Od08g01730 and Od06g06460) of the Rad51 recombinase which plays a role in double strand repair during DNA replication (Lim *et al*., 2020), was upregulated in a similar manner. These observations were consistent with the faster growth rates observed in blue light relative to red and green light and suggested that exposure to blue light signals acceleration of the cell cycle in the RCC809 ecotype of *Ostreococcus*.

The RNA-Seq analysis also revealed changes in expression of genes with key roles in photosynthetic carbon fixation. Three different genes encoding the small subunit of RuBisCO (Od17g01990, Od17g02000 and Od17g02010) showed a 2-fold reduction in expression under blue light relative to red light, with intermediate levels being observed under green light (Fig. S3A-C). RuBisCO activase (Od04g02820), which enhances RuBisCO activity through removal of competitive inhibitors (Portis, 2003), showed similar trends (Fig. S3**d**) suggesting that exposure to blue light may lead to less efficient carbon fixation.

In addition, multiple enzymes in the glycolysis and gluconeogenesis pathways were downregulated under blue light (Fig. S4). While most of these enzymes had both catabolic and anabolic functions, the downregulation of four different fructose-1,6 bisphosphatase genes (Od09g06250, Od06g06980, Od03g02850 and Od20g01100) under blue light suggested a decrease in gluconeogenesis in this condition, while upregulation of pyruvate dehydrogenase suggested an increased rate of glycolysis. Genes associated with fatty acid biosynthesis were also downregulated under blue light (Fig. S5), whereas multiple enzymes in the TCA cycle were upregulated under blue light. This included citrate synthase (Od05g01850), which catalyses the first step of the cycle, and three genes encoding malate dehydrogenase (Od03g03160, Od06g02230, Od08g00070), which regenerates oxaloacetate from malate (Fig. S6). Altogether, these results suggested an increased rate of catabolism and a reduction in energy storage under blue light.

The gene expression data further suggested that synthesis of photopigments (chlorophylls and carotenoids) was globally reduced under blue light, relative to red and green light. 7 genes encoding enzymes in the plastidic MEP/DOXP terpenoid backbone biosynthesis pathway showed reduced expression levels under blue light, with only one showing lower expression under green light (Fig. S7). Multiple enzymes with roles in synthesis of porphyrin precursors of chlorophylls were also downregulated under blue light (Fig. S8**a**), including the enzyme glutamate-1-semialdehyde aminotransferase (GSA-AM) HemL (Od19g01200), which catalyses the production of 5-amino levulinic acid (dALA) from glutamate-1-semialdehyde and is rate-limiting in *Arabidopsis* (Sinha *et al*., 2022).

Expression of chlorophyll *a* (chl*a*) synthase (Od08g01040) was downregulated under blue light (Fig. S8**b**). As this enzyme catalyses the last step of chl*a* biosynthesis, this suggested potential shifts in the ratio of chl*b* to chl*a*. Furthermore, the expression of violaxanthin de-epoxidase (Od11g00170), which catalyses the conversion of violaxanthin to zeaxanthin, was elevated under blue light, whereas zeaxanthin epoxidase (Od02g02570), which catalyses the reverse reaction, was elevated under red light (Fig. S9**a,b**). A member of the CYP97 carotene hydroxylase family (Od09g03950) showed reduced expression under blue light compared to red light (Fig. S9**c**). As this enzyme plays a role in the conversion of carotenes to lutein (Cui *et al*., 2013), this suggested that the synthesis of lutein and its derivatives may decrease under blue light.

Changes related to photosystems were also observed. Two genes with roles in the synthesis of phylloquinone (Od07g04840 and Od20g02670) were downregulated under blue light. Phylloquinone acts as a cofactor in Photosystem I (PSI)-mediated electron transport between the primary electron acceptor and ferredoxin, and a decrease in phylloquinone was previously shown to be associated with a decrease in PSI activity (Gross *et al*., 2006). On the other hand, six genes predicted to encode PSII assembly proteins showed elevated expression under blue light, suggesting an increase in PSII content (Table S2). Expression of another PSII component was reduced suggesting a possible change in composition.

*Ostreococcus* possesses multiple photoreceptors, including a phototropin, a light-oxygen-voltage-histidine kinase (LOV-HK), three cryptochromes, and a histidine kinase rhodopsin (HKR) (Djouani-Tahri *et al*., 2011; Heijde *et al*., 2010; Luck *et al*., 2019; Sullivan *et al*., 2016). While genes involved in light perception were not found to be significantly over-represented within any group of light quality-responsive genes, both genes encoding LOV-HK (Od09g04310 and Od09g04300) showed reduced expression under blue light as compared to red light (Fig. S10). The phototropin photoreceptor (Od15g00300) and the cryptochrome OdCPF1 (Od05g00310) showed a similar response. Expression of the Cry-DASH cryptochrome OdCPF2 (Od01g00210), was elevated in blue light, however. In contrast, that of the histidine kinase rhodopsin (HKR) photoreceptor (Od15g00300) was not affected by light quality.

### Changes in pigment content and photosynthetic parameters in RCC809

To uncover the functional consequences of these changes in gene expression, HPLC analyses of cell pigment contents were carried out. This revealed elevated levels of several carotenoids under blue light relative to red and green light, including neoxanthin, micromonal, dihydrolutein, uriolide and an unknown, previously described carotenoid (Guyon *et al*., 2018). On the other hand, prasinoxanthin, violaxanthin and zeaxanthin levels were reduced (**Fig. 3a**).

**Fig. 3.**
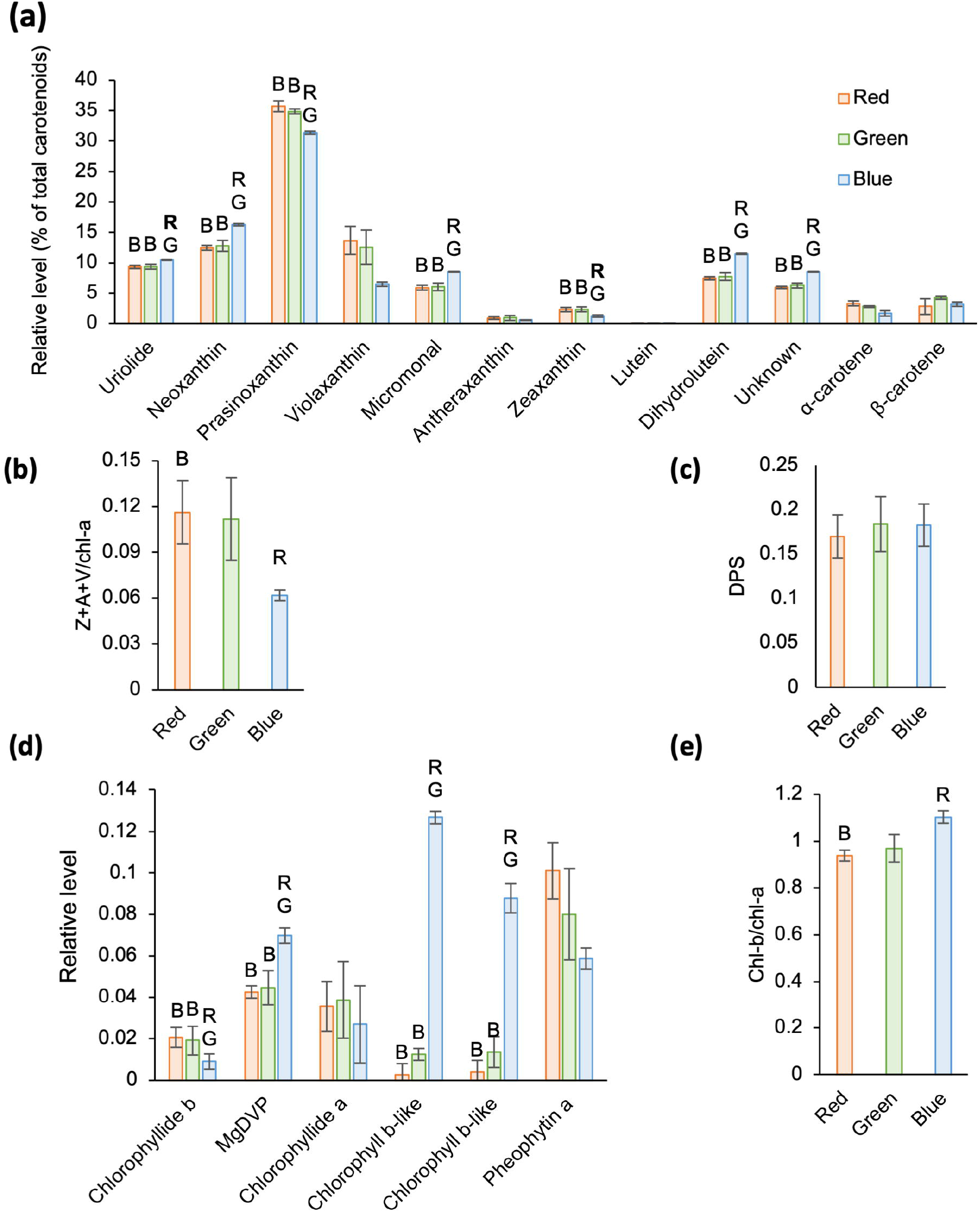
Analysis of pigment contents in RCC809, following acclimation to different light qualities. **(a)** Carotenoid pigments. **(b)** Total xanthophylls relative to chlorophyll *a* (chl*a*). Z, A and V correspond to zeaxanthin, antheraxanthin and violaxanthin, respectively. **(c)** Xanthophyll depoxidation state (DPS), calculated as (Z + 0.5A) / (V + A + Z). **(d)** Chlorophylls (chls) and chlorophyll derivatives. Levels are expressed relative to chl*a*. **(e)** Chl*a* to Chl*b* ratio. Red, green and blue bars correspond to samples acclimated to red, green and blue monochromatic light for 72 hours. Letters indicate significant differences (p < 0.05) with bold letters indicating p < 0.01.

Carotenoids and their oxygenated xanthophyll derivatives play important roles within light-harvesting complexes, both as accessory light-harvesting pigments, and to protect the photosynthetic apparatus from damage by excessive light. The conversion of violaxanthin (V) to antheraxanthin (A) and zeaxanthin (Z) through the de-epoxidation reactions of the xanthophyll cycle acts to protect photosystem II from light damage through quenching of excessive excitation energy and dissipation into heat, a process known as non-photochemical quenching (NPQ) (Goss and Latowski, 2020). The xanthophyll cycle can also play a role in detoxification of reactive oxygen species (ROS) which are generated under supersaturating light conditions. To assess the potential for xanthophyll cycle induction for photoprotection, the total xanthophyll to chl*a* ratio was calculated as Z+A+V/chl*a*, where Z, A and V correspond to relative levels of zeaxanthin, antheraxanthin and violaxanthin, respectively (Demmig-Adams, 1990). This ratio was smaller under blue light than in red or green light, indicating a reduction in the total xanthophyll pool size under blue light relative to red or green light. (**Fig. 3b**). However, no differences were noted in the depoxidation state (DPS), calculated as (Z + 0.5A)/(V + A + Z) (Munné-Bosch and Cela, 2006) (**Fig. 3c**).

As predicted by the gene expression data, blue light induced an increase in chl*b* levels relative to chl*a* (**Fig. 3d,e**). However, the most remarkable response to blue light was an approximately 10-fold increase in two chl*b*-like pigments, relative to red or green light (**Fig. 3d**). The chlorophyll metabolite Mg-2,4-divinyl pheoporphyrin (MgDVP), which plays a role as a light harvesting accessory pigment, also increased significantly relative to chl*a*, almost doubling in blue light compared to both red and green light (**Fig. 3d**). Altogether these results demonstrated that light quality influences the pigment composition of the RCC809 strain of *Ostreococcus*, with potential consequences for light-harvesting efficiency.

To test the consequences of these pigment responses for photosynthetic activity, PAM (pulse-amplitude modulation) fluorometry analysis was used to compare the efficiency of photosystem II after acclimation to the different light qualities. We first assayed the quantum yield of PSII photochemistry (Φ_PSII_) under short pulses of increasing light intensity at 470nm (Ralph and Gademann, 2005). As expected, the quantum yield of PSII was highest in response to low intensity light pulses where carbon fixation operates at maximum photosynthetic efficiency, and decreased at higher light intensities as electron transfer pathways became saturated (**Fig. 4a**). While similar changes were observed in cells adapted to different light conditions, Φ_PSII_ was always highest in cells acclimated to red light and lowest in cells acclimated to blue light at a given intensity of excitatory light.

**Fig. 4.**
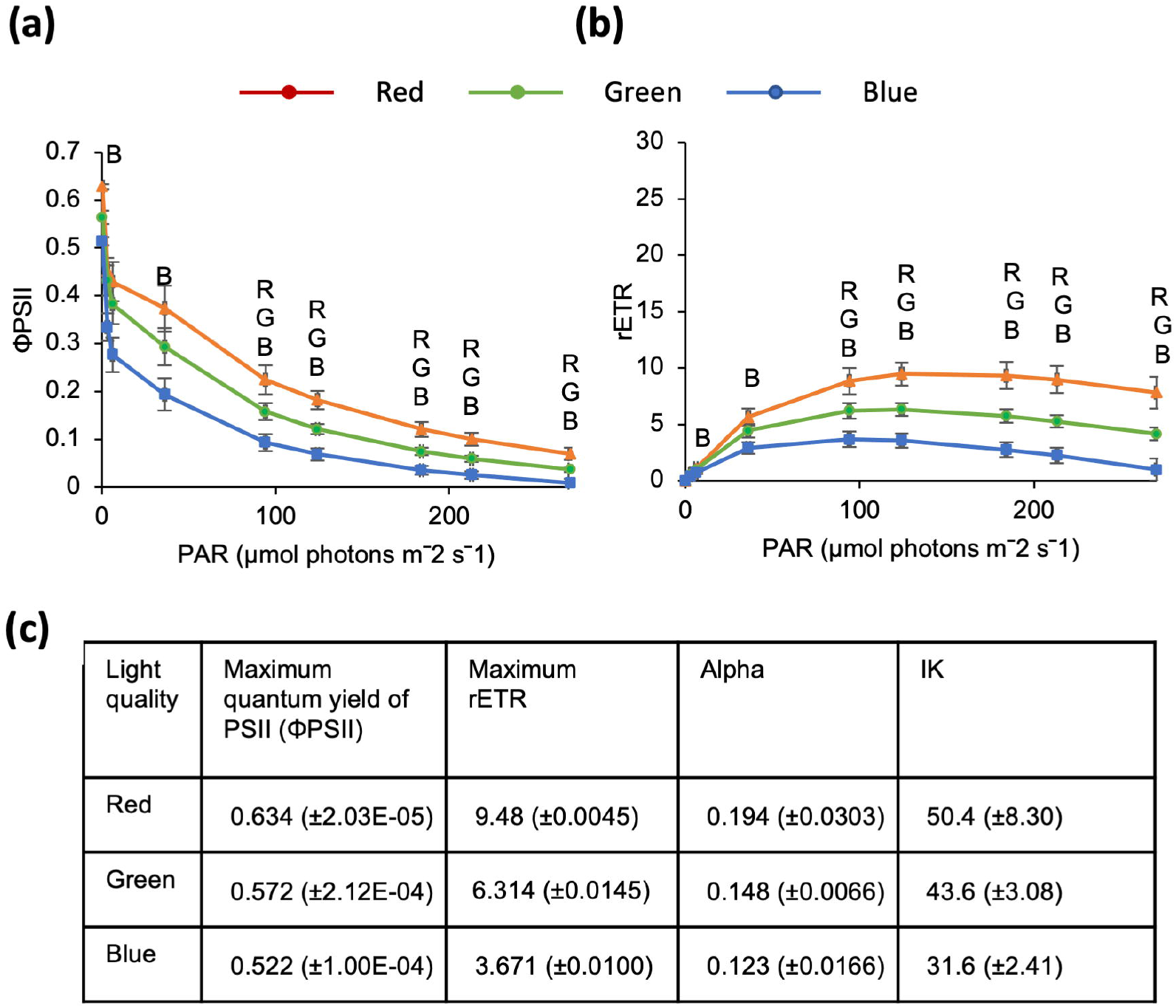
Effect of light quality on photosynthetic parameters in RCC8009. **(a)** and **(b)** Photosynthetic light-response curves showing the quantum yield of Photosystem II (Φ SII) and relative electron transport rate (rETR) as a function of irradiance (photosynthetically active radiation or PAR), for cultures acclimated to either red, green or blue light. Letters indicate significant differences (p < 0.05) with bold letters indicating p < 0.01. **(c)** Maximum quantum yield, rETR Max, Alpha and IK values.

The relative electron transport rate (rETR) increased with the intensity of excitation, reaching a plateau around 50-150μmol photons m^-2^ s^-1^, then decreased at higher light intensities (**Fig. 4b**). However, higher electron transfer rates were observed in cells acclimated to red light, and lower rates in cells acclimated to blue light. Furthermore, saturation was reached at a lower intensity in cells acclimated to blue light (about 50μmol photons m^-2^ s^-1^ as compared to 150μmol photons m^-2^ s^-1^ in cells acclimated to red or green light).

These findings suggest that the photosynthetic machinery in RCC809 tolerates higher light intensities after acclimation to red light as compared to blue light. Green light acclimated cultures showed intermediate properties.

### Light quality responses are attenuated in OTTH0595, a shallow water ecotype of *Ostreococcus*

While light intensity is thought to be the primary abiotic factor determining the clade distribution of *Ostreococcus* species (Botebol *et al*., 2017; Demir-Hilton *et al*., 2011; Rodriguez *et al*., 2005; Six *et al*., 2009), we hypothesized that light quality may also play a role. Our prediction was that an ecotype originating from a shallow water environment and exposed to the full visible light spectrum at all times may exhibit distinct responses to light quality from a deep ocean ecotype, which is exposed to variable wavelengths of light as it moves up and down the water column. We therefore compared the light quality responses of the deep ocean ecotype, RCC809 with those of a lagoon species, OTTH0595, also known as *O. tauri*.

OTTH0595 growth rates were only slightly slower under red light compared to blue and green light (**Fig. 5a**). HPLC analyses revealed that the chl*b*:chl*a* rati*o* was not affected by light quality (**Fig. 5b**). Small increases in micromonal and dihydrolutein contents were observed in blue light (9% and 14 %, respectively; **Fig. 5c**), but these were minor changes compared to those observed in RCC809 (43% and 52%, respectively). Consistent with previous observations that OTTH0595 tolerates higher light intensities (Cardol *et al*., 2008; Demir-Hilton *et al*., 2011; Six *et al*., 2008; Six *et al*., 2009), rETR values were higher than in RCC809 and did not decrease at high light intensities. No significant differences in PSII quantum yield or rETR were observed between light qualities, however (**Fig. 5d,e**).

**Fig. 5.**
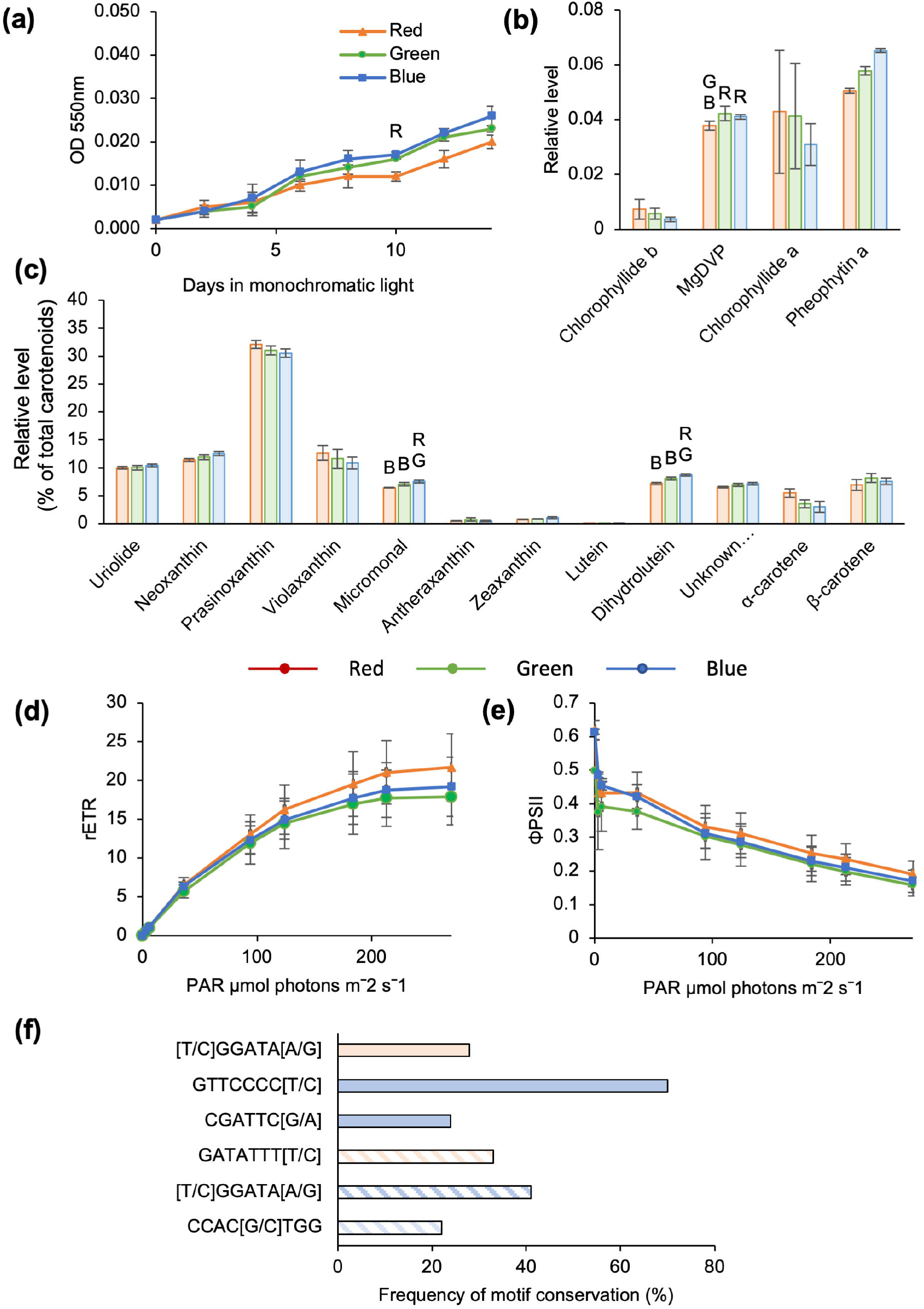
Light quality responses in the lagoon species, OTTH0595, also known as *tauri*. **(a)** Growth kinetics. **(b) and (c)** Levels of chls and chl derivatives (relative to chla), and levels of carotenoid pigments (relative to total carotenoids). Red, green and blue bars correspond to samples acclimated to red, green and blue monochromatic light for 72 hours. Letters indicate significant differences (p < 0.05). **(d) and (e)** Photosynthetic light-response curves showing the quantum yield of Photosystem II (ΦPSII) and relative electron transport rate (rETR) as a function of irradiance (photosynthetically active radiation or PAR), for cultures acclimated to either red, green or blue light. **(f)** Frequency of motif conservation between RCC809 red and blue light-responsive genes and their OTTH0595 homologues. Solid red and blue bars correspond to genes upregulated under red and blue light, respectively. Hatched red and blue bars correspond to genes downregulated under red and blue light.

To examine whether the attenuated light responses of the OTTH0595 species were linked to divergence in gene regulation relative to RCC809, we identified the OTTH0595 homologues of light quality-responsive genes from RCC809 and examined their upstream regulatory sequences for the presence or absence of the candidate regulatory elements identified in Table 2. While the GTTCCCC[T/C] motif, recognised by the cell cycle-related transcription factor E2F was conserved in 80% of OTTH0595 homologues, other motifs were only present in 30 to 40% of OTTH0595 genes, suggesting that they had been lost during the course of its evolutionary divergence from RCC809 (**Fig. 5f**).

## Discussion

### Blue light signals a distinct growth strategy from red and green light in RCC809

Our results demonstrate that the *Ostreococcus* ecotype RCC809 exhibits responses to light quality. Transcriptomic analyses revealed differences in gene expression between cells acclimated to red, green, and blue monochromatic light. Gene expression under blue light was most distinct from red light, whilst intermediate phenotypes were seen under green light. Blue light signalled a switch in physiology and metabolism to a state that favoured growth at the expense of energy storage. Thus, acclimation to blue light resulted in the elevated expression of genes associated with the cell cycle and faster growth rates. Conversely, expression of enzymes involved in starch or fatty acid biosynthesis was reduced, whereas the expression of TCA cycle enzymes was elevated. Exposure to green or red light elicited the opposite response, suggesting that enrichment for red or green wavelengths may signal conditions less favourable for growth in the natural environment, such as shading by sediments or other particulate material, competition from other organisms, or exposure to damaging light intensities near the surface.

We questioned whether the gene expression signatures observed under green light might represent a starvation response due to poor absorption of light by chlorophylls in this part of the spectrum. However, the fact that similar responses were observed under red and green light argued against this, because red light is absorbed efficiently by chlorophylls (Liu and Van Lersel, 2021). Furthermore, the growth arrest observed under red or green light was accompanied by upregulation of starch and fatty acid biosynthesis pathways, which is not consistent with a starvation response.

### RCC809 adjusts its pigment contents and photosystem function as a function of the light quality environment

Previous work showed that increases in light intensity induce the downregulation of both chl*a* and chl*b* content, accumulation of lutein, and increased xanthophyll de-epoxidation (Six *et al*., 2008). Blue light triggered a distinct set of responses in our experiment, suggesting that cells did not respond to the higher energy levels carried by blue wavelengths, but rather to the specific light quality.

Blue light induced an increase in the carotenoid pigments, dihydrolutein and micromonal, as well as an increase in MgDVP, chl*b* and chl*b*-like pigments relative to chl*a*. This was consistent with chromatic adaptation, as chl*b* and MgDVP absorb much more under blue light than chl*a* and these changes would be expected to enhance cells ability to utilise blue light for photosynthesis (Raven, 1996).

In contrast, red and green light induced expression of enzymes involved in the biosynthesis of chl*a*, which would be predicted to increase light absorption under red light. Whilst the lower chl*b*:chl*a* ratio suggested that the size of the photosynthetic antenna was smaller in cells acclimated to red light, PSII efficiency was increased as indicated by higher PSII quantum yield and rETR values. This may be linked to higher RuBisCO activity, as the expression of RuBisCO small subunits and of RuBisCO activase was higher under red and green light, compared to blue light, and the RuBisCo activation state is known to be positively correlated with the electron transport rate in higher plants (Perdomo *et al*., 2017).

An increase in the total xanthophyll pool size was also observed under red and green light, indicating an increased potential for photoprotection. Consistent with this observation, red or green light-adapted cultures maintained higher PSII quantum yields and rETR under increasing light intensities. Detection of green and red light may allow cells to perceive their proximity to the surface and to anticipate exposure to high light intensities through induction of photoprotective mechanisms.

### Mechanism of light quality responses

The response of RCC809 to changes in light quality contrasts with that known for land plants, where the effects of green light oppose those of red and blue wavelengths (Wang and Folta, 2013). This distinct pattern of light quality responses may be due to the presence of a distinct set of photoreceptors. In *Ostreococcus*, the cryptochrome, LOV-HK and phototropin photoreceptors, which absorb specifically in the UV/blue light range, may allow specific responses to blue light, whereas green light may be perceived by HKR, which absorbs maximally at 505nm in the dark state or 560nm in the presence of salt (Luck *et al*., 2019). The identity of the photoreceptor responsible for perception of red light remains unclear. It is possible that no such photoreceptor is present, and that *Ostreococcus* simply responds to the presence or absence of blue light.

### Evidence for adaptation to different light quality environments

We compared the light quality responses of the RCC809 ecotype, sourced from the open Atlantic Ocean at a depth of 105 metres, with those of the lagoon ecotype OTTH0595. These two Ostreococcus species were previously shown to belong to distinct clades (Demir-Hilton *et al*., 2011) and to exhibit distinct responses to light intensity (Cardol *et al*., 2008; Six *et al*., 2008; Six *et al*., 2009). OTTH0595 originates from shallow environments where it is not greatly affected by the narrowing of spectral availability associated with depth (Potes *et al*., 2013). Instead, the spectral quality varies mostly according to the level of suspended sediment and other photosynthetic organisms. It may therefore be less important for this species to be able to perceive the blue-light enriched environment associated with depth, and to adjust its photosystems accordingly. Furthermore, OTTH0595 showed no differences in pigment contents when adapted to different light qualities, except for a small increase in dihydrolutein and micromonal under blue light, and its PSII quantum yield and rETR were unaffected.

While OTTH0595 possesses the same set of photoreceptors as RCC809, mechanistic clues to their distinct responses to light quality may be obtained through investigation of promoter motifs and their cognate transcription factors. Analysis of over-represented motifs within RCC809 light quality-responsive promoters identified a number of candidate regulatory motifs, including binding sites for MYB and bHLH transcription factors. Remarkably, most of these promoter motifs were missing from OTTH0595 homologues of RCC809 light quality-responsive genes, suggesting a lack of selective pressure for such regulation in OTTH0595. Furthermore, a single bHLH transcription factor was identified in OTTH0595, as compared to four in RCC809, suggesting the loss of light-responsive transcription factors.

Taken altogether, these observations suggest that whilst light intensity has been considered the primary abiotic factor determining the distribution of *Ostreococcus* ecotypes, differences in light quality responses of ecotypes should also be considered as an important factor in determining adaptation to distinct environmental niches.

## Supplementary Information

*Fig. S1.* Spectroradiometer analysis of the three light conditions used in our experiments.

*Fig. S2.* Examples of cell cycle-related genes upregulated under blue light.

*Fig. S3.* Effects of light quality on the expression of Calvin cycle-related genes.

*Fig. S4.* Effects of light quality on the expression of glycolysis and starch synthesis-related genes.

*Fig. S5.* Effects of light quality on the expression of fatty acid synthesis-related genes.

*Fig. S6.* Effects of light quality on the expression of TCA cycle genes.

*Fig. S7.* Effects of light quality on the expression of terpenoid biosynthesis genes.

*Fig. S8.* Effects of light quality on the expression of genes involved in porphyrin and chlorophyll metabolism.

*Fig. S9.* Effects of light quality on the expression of carotenoid biosynthesis genes.

*Fig. S10.* Effects of light quality on the expression of *Ostreococcus* RCC809 genes encoding photoreceptors.

*Table S1.* Lists of differentially expressed genes identified for RCC809 in pairwise comparisons between red and green, red and blue, or blue and green light conditions.

*Table S2.* Light quality-responsive genes related to the cell cycle.

*Table S3.* Light quality-responsive genes with roles in photosystem II assembly.

## Acknowledgements

The RCC809 and OTTH0595 cell lines were obtained from the Roscoff culture collection. Sequencing of RNA-Seq samples was carried out at the Wellcome Trust, and RNA-Seq data were analysed within the Cyverse Discovery Environment (https://cyverse.org).

## Author contributions

SD carried out prepared the samples used in RNA-Seq analyses. ES analysed the RNA-Seq data, performed growth, pigment and PhytoPAM analyses with the help of RP. DS and IAC conceived and supervised the project. FYB assisted with preliminary experiments. ES and IAC prepared the manuscript.

## Data availability

The transcriptomics dataset was deposited on the Gene Expression Omnibus database under the accession number GSE221420.

## References

Andrews S. 2015. FastQC: A Quality Control Tool for High Throughput Sequence Data [Online] Available Online at http://www.bioinformatics.babraham.ac.uk/projects/fastqc/.

Bailey TL. 2011. DREME: motif discovery in transcription factor ChIP-seq data. Bioinformatics 27, 1653–1659.

Blanc-Mathieu R, Verhelst B, Derelle E, Rombauts S, Bouget F, Carre I, Chateau A, Eyre-Walker A, Grimsley N, Moreau H, Piegu B, Rivals E, Schackwitz W, Van de Peer Y, Piganeau G. 2014. An improved genome of the model marine alga Ostreococcus tauri unfolds by assessing Illumina de novo assemblies. BioMed Central Genomics 15,1103.

Bolger AM, Lohse M, Usadel B. 2014. Trimmomatic: a flexible trimmer for Illumina sequence data. Bioinformatics 30, 2114–2120.

Botebol H, Lelandais G, Six C, Lesuisse E, Meng A, Bittner L, Lecrom S, Sutak R, Lozano JC, Schatt P, Vergé V, Blain S, Bouget FY. 2017. Acclimation of a low iron adapted Ostreococcus strain to iron limitation through cell biomass lowering. Scientific Reports 7, 327.

Bouman HA, Platt T, Kraay GW, Sathyendranath S, Irwin BD. 2000. Bio-optical properties of the subtropical North Atlantic. I. Vertical variability. Marine Ecology Progress Series 200, 3–18.

Brunet C, Chandrasekaran R, Barra L, Giovagnetti V, Corato F, Ruban AV. 2014. Spectral Radiation Dependent Photoprotective Mechanism in the Diatom Pseudo-nitzschia multistriata. Plos One 9, e87015.

Camacho C, Coulouris G, Avagyan V, Ma N, Papadopoulos J, Bealer K, Madden TL. 2009. BLAST+: architecture and applications. BioMed Central Bioinformatics 10, 421.

Cardol P, Bailleul B, Rappaport F, Derelle E, Beal D, Breyton C, Bailey S, Wollman F, Grossman A, Moreau H, Finazzi G. 2008. An original adaptation of photosynthesis in the marine green alga *Ostreococcus*. Proceedings of the National Academy of Sciences of the United States of America 105, 7881–7886.

Chretiennot-Dinet M, Courties C, Vaquer A, Neveux J, Claustre H, Lautier J, Machado M. 1995. A new marine picoeucaryote – *Ostreococcus tauri* gen et sp-nov (Chlorophyta, Prasinophyceae). Phycologia 34, 285–292.

Corellou F, Camasses A, Ligat L, Peaucellier G, Bouget F. 2005. Atypical regulation of a green lineage-specific B-type cyclin-dependent kinase. Plant Physiology 138, 1627–1636.

Corellou F, Schwartz C, Motta J, Djouani-Tahri E, Sanchez F, Bouget F. 2009. Clocks in the green lineage: comparative functional analysis of the circadian architecture of the Picoeukaryote *Ostreococcus*. Plant Cell 21, 3436–3449.

Cui H, Yu X, Wang Y, Cui Y, Li X, Liu Z, Qin S. 2013. Evolutionary origins, molecular cloning and expression of carotenoid hydroxylases in eukaryotic photosynthetic algae. BioMed Central Genomics 14, 457.

Demir-Hilton E, Sudek S, Cuvelier M, Gentemann C, Zehr J, Worden A. 2011. Global distribution patterns of distinct clades of the photosynthetic picoeukaryote *Ostreococcus*. ISME Journal 5, 1095–1107.

Demmig-Adams B. 1990. Carotenoids and photoprotection in plants: A role for the xanthophyll zeaxanthin. Biochimica et Biophysica Acta 1020, 1–24.

Derelle E, Ferraz C, Rombauts S, Rouze P, Worden A, Robbens S, Partensky F, Degroeve S, Echeynie S, Cooke R, Saeys Y, Wuyts J, Jabbari K, Bowler C, Panaud O, Piegu B, Ball S, Ral J, Bouget F, Piganeau G, De Baets B, Picard A, Delseny M, Demaille J, Van de Peer Y, Moreau H. 2006. Genome analysis of the smallest free-living eukaryote *Ostreococcus tauri* unveils many unique features. Proceedings of the National Academy of Sciences of the United States of America 103, 11647–11652.

Djouani-Tahri E-B, Christie JM, Sanchez-Ferandin S, Sanchez F, Bouget F-Y, Corellou F. 2011. A eukaryotic LOV-histidine kinase with circadian clock function in the picoalga Ostreococcus. The Plant Journal 65, 578–588.

Erga S, Ssebiyonga N, Frette O, Hamre B, Aure J, Strand O, Strohmeier T. 2012. Dynamics of phytoplankton distribution and photosynthetic capacity in a western Norwegian fjord during coastal upwelling: Effects on optical properties. Estuarine Coastal and Shelf Science 97, 91–103.

Farinas B, Mary C, de O Manes CL, Bhaud Y, Peaucellier G, Moreau H. 2006. Natural synchronisation for the study of cell division in the green unicellular alga *Ostreococcus tauri*. Plant Molecular Biology 60, 277–292.

Franklin KA, Whitelam GC. 2005. Phytochromes and Shade-avoidance Responses in Plants. Annals of Botany 96, 169–175.

Goff L, Trapnell C, Kelley D. 2020. Analysis, exploration, manipulation, and visualization of Cufflinks high-throughput sequencing data. R package version 2.40.0.

Goss R, Latowski D. 2020. Lipid Dependence of Xanthophyll Cycling in Higher Plants and Algae. Frontiers in plant science 11, 455.

Götz S, García-Gómez JM, Terol J, Williams TD, Nagaraj SH, Nueda MJ, Robles M, Talón M, Dopazo J, Conesa A. 2008. High-throughput functional annotation and data mining with the Blast2GO suite. Nucleic Acids Research 36, 3420–3435.

Grigoriev IV, Nordberg H, Shabalov I, Aerts A, Cantor M, Goodstein D, Kuo A, Minovitsky S, Nikitin R, Ohm RA, Otillar R, Poliakov A, Ratnere I, Riley R, Smirnova T, Rokhsar D, Dubchak I. 2012. The genome portal of the Department of Energy Joint Genome Institute. Nucleic Acids Research 40, D26–32.

Gross J, Cho WK, Lezhneva L, Falk J, Krupinska K, Shinozaki K, Seki M, Herrmann RG, Meurer J. 2006. A plant locus essential for phylloquinone (vitamin K1) biosynthesis originated from a fusion of four eubacterial genes. J Biol Chem 281, 17189–17196.

Gruening BA. 2014. Galaxy Wrapper [ONLINE] https://github.com/bgruening/galaxytools.

Guyon JB, Vergé V, Schatt P, Lozano JC, Liennard M, Bouget FY. 2018. Comparative Analysis of Culture Conditions for the Optimization of Carotenoid Production in Several Strains of the Picoeukaryote Ostreococcus. Mar Drugs 16, 76.

Heijde M, Zabulon G, Corellou F, Ishikawa T, Brazard J, Usman A, Sanchez F, Plaza P, Martin M, Falciatore A, Todo T, Bouget F-Y, Bowler C. 2010. Characterization of two members of the cryptochrome/photolyase family from Ostreococcus tauri provides insights into the origin and evolution of cryptochromes. Plant, Cell & Environment 33, 1614–1626.

Hintz NH, Zeising M, Striebel M. 2021. Changes in spectral quality of underwater light alter phytoplankton community composition. Limnology and Oceanography 66, 3327–3337.

Jardillier L, Zubkov M, Pearman J, Scanlan D. 2010. Significant CO_2_ fixation by small prymnesiophytes in the subtropical and tropical northeast Atlantic Ocean. The ISME Journal: Multidisciplinary Journal of Microbial Ecology 4, 1180–1192.

Keller MD, Selvin RC, Claus W, Guillard RRL. 1987. Media for the culture of oceanic ultraphytoplankton. Journal of Phycology 23, 633–638.

Kim D, Pertea G, Trapnell C, Pimentel H, Kelley R, Salzberg SL. 2013. TopHat2: accurate alignment of transcriptomes in the presence of insertions, deletions and gene fusions. Genome Biology 14, R36.

Kirkham AR, Lepère C, Jardillier LE, Not F, Bouman H, Mead A, Scanlan DJ. 2013. A global perspective on marine photosynthetic picoeukaryote community structure. The ISME Journal: Multidisciplinary Journal of Microbial Ecology 7, 922–936.

Kolde R. 2019. pheatmap: Pretty Heatmaps. R package version 1.0.12.

Kulk G, Platt T, Dingle J, Jackson T, Jönsson BF, Bouman HA, Babin M, Brewin RJW, Doblin M, Estrada M, Figueiras FG, Furuya K, González-Benítez N, Gudfinnsson HG, Gudmundsson K, Huang B, Isada T, Kovač Ž, Lutz VA, Marañón E, Raman M, Richardson K, Rozema PD, Poll WHvd, Segura V, Tilstone GH, Uitz J, Dongen-Vogels Vv, Yoshikawa T, Sathyendranath S. 2020. Primary Production, an Index of Climate Change in the Ocean: Satellite-Based Estimates over Two Decades. Remote Sensing 12, 826.

Kume A, Akitsu T, Nasahara KN. 2018. Why is chlorophyll b only used in light-harvesting systems? Journal of Plant Research 131, 961–972.

Langmead B, Salzberg SL. 2012. Fast gapped-read alignment with Bowtie 2. Nature Methods 9, 357–359.

Li Y, Cai X, Gu W, Wang G. 2020. Transcriptome analysis of carotenoid biosynthesis in Dunaliella salina under red and blue light. Journal of Oceanology and Limnology 38, 177–185.

Lichtenthaler HK. 1987. [34] Chlorophylls and carotenoids: Pigments of photosynthetic biomembranes. Methods in Enzymology, Vol. 148: Academic Press, 350–382.

Lim G, Chang Y, Huh W-K. 2020. Phosphoregulation of Rad51/Rad52 by CDK1 functions as a molecular switch for cell cycle–specific activation of homologous recombination. Science Advances 6, eaay2669.

Limardo AJ, Sudek S, Choi CJ, Poirier C, Rii YM, Blum M, Roth R, Goodenough U, Church MJ, Worden AZ. 2017. Quantitative biogeography of picoprasinophytes establishes ecotype distributions and significant contributions to marine phytoplankton. Environmental Microbiology 19, 3219–3234.

Liu J, Van Lersel MW. 2021. Photosynthetic Physiology of Blue, Green, and Red Light: Light Intensity Effects and Underlying Mechanisms. Frontiers in plant science 12, 619987.

Luck M, Velázquez Escobar F, Glass K, Sabotke M-I, Hagedorn R, Corellou F, Siebert F, Hildebrandt P, Hegemann P. 2019. Photoreactions of the Histidine Kinase Rhodopsin Ot-HKR from the Marine Picoalga Ostreococcus tauri. Biochemistry 58, 1878–1891.

Maere S, Heymans K, Kuiper M. 2005. BiNGO: a Cytoscape plugin to assess overrepresentation of gene ontology categories in biological networks. Bioinformatics 21, 3448–3449.

Mullaney PF, Van Dilla MA, Coulter JR, Dean PN. 1969. Cell sizing: a light scattering photometer for rapid volume determination. Review of Scientific Instruments 40, 1029–1032.

Mullin MM. 2001. Plankton. In: Cochran JK, Bokuniewicz HJ, Yager PL, eds. Encyclopedia of Ocean Sciences (Third Edition). Oxford: Academic Press, 613–614.

Munné-Bosch S, Cela J. 2006. Effects of water deficit on photosystem II photochemistry and photoprotection during acclimation of lyreleaf sage (*Salvia lyrata L*.) plants to high light. Journal of Photochemistry and Photobiology B-Biology 85, 191–197.

Neun S, Hintz NH, Schröder M, Striebel M. 2022. Phytoplankton Response to Different Light Colors and Fluctuation Frequencies. Frontiers in Marine Science 9, 824624.

Palenik B, Grimwood J, Aerts A, Rouze P, Salamov A, Putnam N, Dupont C, Jorgensen R, Derelle E, Rombauts S, Zhou K, Otillar R, Merchant S, Podell S, Gaasterland T, Napoli C, Gendler K, Manuell A, Tai V, Vallon O, Piganeau G, Jancek S, Heijde M, Jabbari K, Bowler C, Lohr M, Robbens S, Werner G, Dubchak I, Pazour G, Ren Q, Paulsen I, Delwiche C, Schmutz J, Rokhsar D, Van de Peer Y, Moreau H, Grigoriev I. 2007. The tiny eukaryote *Ostreococcus* provides genomic insights into the paradox of plankton speciation. Proceedings of the National Academy of Sciences of the United States of America 104, 7705–7710.

Patelou M, Infante C, Dardelle F, Randewig D, Kouri ED, Udvardi MK, Tsiplakou E, Mantecón L, Flemetakis E. 2020. Transcriptomic and metabolomic adaptation of Nannochloropsis gaditana grown under different light regimes. Algal Research 45, 101735.

Perdomo JA, Capó-Bauçà S, Carmo-Silva E, Galmés J. 2017. Rubisco and Rubisco Activase Play an Important Role in the Biochemical Limitations of Photosynthesis in Rice, Wheat, and Maize under High Temperature and Water Deficit. Frontiers in plant science 8, 490.

Platt T, Gallegos CL, Harrison WG. 1980. PHOTOINHIBITION OF PHOTOSYNTHESIS IN NATURAL ASSEMBLAGES OF MARINE PHYTOPLANKTON. Journal of Marine Research 38, 687–701.

Portis AR, Jr. 2003. Rubisco activase – Rubisco’s catalytic chaperone. Photosynthesis Research 75, 11–27.

Potes M, Costa MJ, Salgado R, Bortoli D, Serafim A, Moigne PL. 2013. Spectral measurements of underwater downwelling radiance of inland water bodies. Tellus A: Dynamic Meteorology and Oceanography 65, 20774.

Quinlan AR, Hall IM. 2010. BEDTools: a flexible suite of utilities for comparing genomic features. Bioinformatics 26, 841–842.

Ralph PJ, Gademann R. 2005. Rapid light curves: A powerful tool to assess photosynthetic activity. Aquatic Botany 82, 222–237.

Raven JA. 1996. The Bigger The Fewer: Size, Taxonomic Diversity and The Range of Chlorophyll(Ide) Pigments in Oxygen-Evolving Marine Photolithotrophs. Journal of the Marine Biological Association of the United Kingdom 76, 211–217.

Rii YM, Duhamel S, Bidigare RR, Karl DM, Repeta DJ, Church MJ. 2016. Diversity and productivity of photosynthetic picoeukaryotes in biogeochemically distinct regions of the South East Pacific Ocean. Limnology and Oceanography 61, 806–824.

Rodriguez F, Derelle E, Guillou L, Le Gall F, Vaulot D, Moreau H. 2005. Ecotype diversity in the marine picoeukaryote *Ostreococcus* (Chlorophyta, Prasinophyceae). Environmental Microbiology 7, 853–859.

Schaum CE, Rost B, Collins S. 2016. Environmental stability affects phenotypic evolution in a globally distributed marine picoplankton. The Isme Journal 10, 75–84.

Schubert H, Sagert S, Forster R. 2001. Evaluation of the different levels of variability in the underwater light field of a shallow estuary. Helgoland Marine Research 55, 12–22.

Sinha N, Eirich J, Finkemeier I, Grimm B. 2022. Glutamate 1-semialdehyde aminotransferase is connected to GluTR by GluTR-binding protein and contributes to the rate-limiting step of 5-aminolevulinic acid synthesis. Plant Cell 34, 4623–4640.

Six C, Finkel Z, Rodriguez F, Marie D, Partensky F, Campbell D. 2008. Contrasting photoacclimation costs in ecotypes of the marine eukaryotic picoplankter *Ostreococcus*. Limnology and Oceanography 53, 255–265.

Six C, Sherrard R, Lionard M, Roy S, Campbell D. 2009. Photosystem II and pigment dynamics among ecotypes of the green alga *Ostreococcus*. Plant Physiology 151, 379–390.

Sullivan S, Petersen J, Blackwood L, Papanatsiou M, Christie JM. 2016. Functional characterization of Ostreococcus tauri phototropin. New Phytologist 209, 612–623.

Sun D, Li Y, Wang Q, Lv H, Le C, Huang C, Gong S. 2010. Partitioning particulate scattering and absorption into contributions of phytoplankton and non-algal particles in winter in Lake Taihu (China). Hydrobiologia 644, 337–349.

The_Galaxy_Community. 2022. The Galaxy platform for accessible, reproducible and collaborative biomedical analyses: 2022 update. Nucleic Acids Research 50, W345–W351.

Tilzer M, Stambler N, Lovengreen C. 1995. The role of phytoplankton in determining the underwater light climate in Lake Constance. Hydrobiologia 316, 161–172.

Trapnell C, Hendrickson DG, Sauvageau M, Goff L, Rinn JL, Pachter L. 2013. Differential analysis of gene regulation at transcript resolution with RNA-seq. Nature Biotechnology 31, 46–53.

Van Heukelem L, Thomas CS. 2001. Computer-assisted high-performance liquid chromatography method development with applications to the isolation and analysis of phytoplankton pigments. The Journal of Chromatography A 910, 31–49.

Wang Y, Folta KM. 2013. Contributions of green light to plant growth and development. American journal of botany 100, 70–78.

Weirauch MT, Yang A, Albu M, Cote AG, Montenegro-Montero A, Drewe P, Najafabadi HS, Lambert SA, Mann I, Cook K, Zheng H, Goity A, van Bakel H, Lozano JC, Galli M, Lewsey MG, Huang E, Mukherjee T, Chen X, Reece-Hoyes JS, Govindarajan S, Shaulsky G, Walhout AJM, Bouget FY, Ratsch G, Larrondo LF, Ecker JR, Hughes TR. 2014. Determination and inference of eukaryotic transcription factor sequence specificity. Cell 158, 1431–1443.

Xu L, Pan W, Yang G, Tang X, Martin RM, Liu G, Zhong C. 2021. Impact of light quality on freshwater phytoplankton community in outdoor mesocosms. Environmental Science and Pollution Research 28, 58536–58548.

Yuan H, Zhang X, Jiang Z, Wang X, Wang Y, Cao L, Zhang X. 2020. Effect of light spectra on microalgal biofilm: Cell growth, photosynthetic property, and main organic composition. Renewable Energy 157, 83–89.

